# Warming winter disrupts mycorrhizal phenology and plant-fungal nutrient cycling

**DOI:** 10.64898/2025.12.11.693722

**Authors:** Hannah B Shulman, Aimée T Classen, Ian Breckheimer, Wenming Dong, Peter Falb, Amanda Henderson, David W Inouye, Eun Ji Shin, Patrick Sorensen, Olivia K Vought, Stephanie N Kivlin

## Abstract

Climate change is reshaping the timing of ecological processes in montane ecosystems, where short, snowpack-dependent growing seasons tightly constrain plant–microbe interactions. Arbuscular mycorrhizal (AM) fungi regulate plant acquisition of nitrogen (N) and phosphorus (P) during nutrient pulses triggered by melting snowpack. Yet the extent and consequences of warming-induced phenological asynchrony between AM fungi and host plants remains unknown. We experimentally advanced snowmelt in a subalpine meadow and monitored plant and AM fungal growth, soil nutrients, and AM fungal community composition throughout the growing season. Early snowmelt advanced plant greenness and root standing stock but suppressed AM fungal hyphal production, reducing available NH□□ and PO_4_^3-^. Hyphal allocation strategies shaped AM fungal temporal niches and species-specific responses to warming. Nutrient-foraging, edaphophilic AM fungi dominated early in the season, while rhizophilic AM fungi dominated later, following peak root growth. Our results reveal that fungal functional traits and nutrient dynamics govern the temporal niche partitioning of mycorrhizal fungi. By decoupling AM fungal activity from plant demand and nutrient mineralization windows, early snowmelt drives plant-fungal asynchrony. This decoupling weakens mycorrhizal symbioses and threatens nutrient retention and ecosystem stability under future warming. Such belowground temporal mismatches may determine the resilience of seasonally temperature-dependent ecosystems under warming.

## Introduction

In montane and poleward ecosystems, plant phenology and nutrient cycling are tightly regulated by the temporal constraints imposed by long winters^1^. Within this narrow growing season window, mycorrhizal fungal associations play a central role in mediating plant access to nitrogen (N) and phosphorus (P)^2^. Earlier snowmelt, altered precipitation regimes, and warming temperatures are shifting the phenologies of plants and soil microbes, leading to increased potential for mismatches in growth and nutrient demand ^3,4^. Disruption of mycorrhizal interactions through phenological asynchrony may feed back to the rate of climate change by altering nutrient pools and fluxes at ecosystem scales, potentially leading to nutrient loss from terrestrial ecosystems^5^. In this study, we investigated the mycorrhizal response to early snowmelt and the consequences on soil nutrient cycling, to determine if phenological asynchrony between mycorrhizal fungi and host plants alters ecosystem function as it does with other mutualists^6^.

Where the growing season is limited by cold winters and deep snow, a significant amount of biogeochemical processes that provision plant nutrients occur during the snowmelt period, which begins when meltwater reaches the soil surface and lasts until the snowpack has disappeared^7^. The role of arbuscular mycorrhizal (AM) fungi during snowmelt is unclear, although there is compelling evidence that mycorrhizas are key to early-season nutrient acquisition: fungal colonization of alpine plant roots may persist through winter in order to absorb the post-snowmelt nutrient pulse^8^. Biotic and abiotic shifts in the soil, such as microbial biomass and NH_4_^+^ accumulation under snow, may cue mycorrhizal phenology and drive major changes in nutrient mineralization and availability ^9^ Disruptions to plant–fungal synchrony during this short snowmelt window (Fig 1A) may be critical, as nutrients released by microbial turnover are a key source of N and P for plants^10,11^.

**Figure 1:**
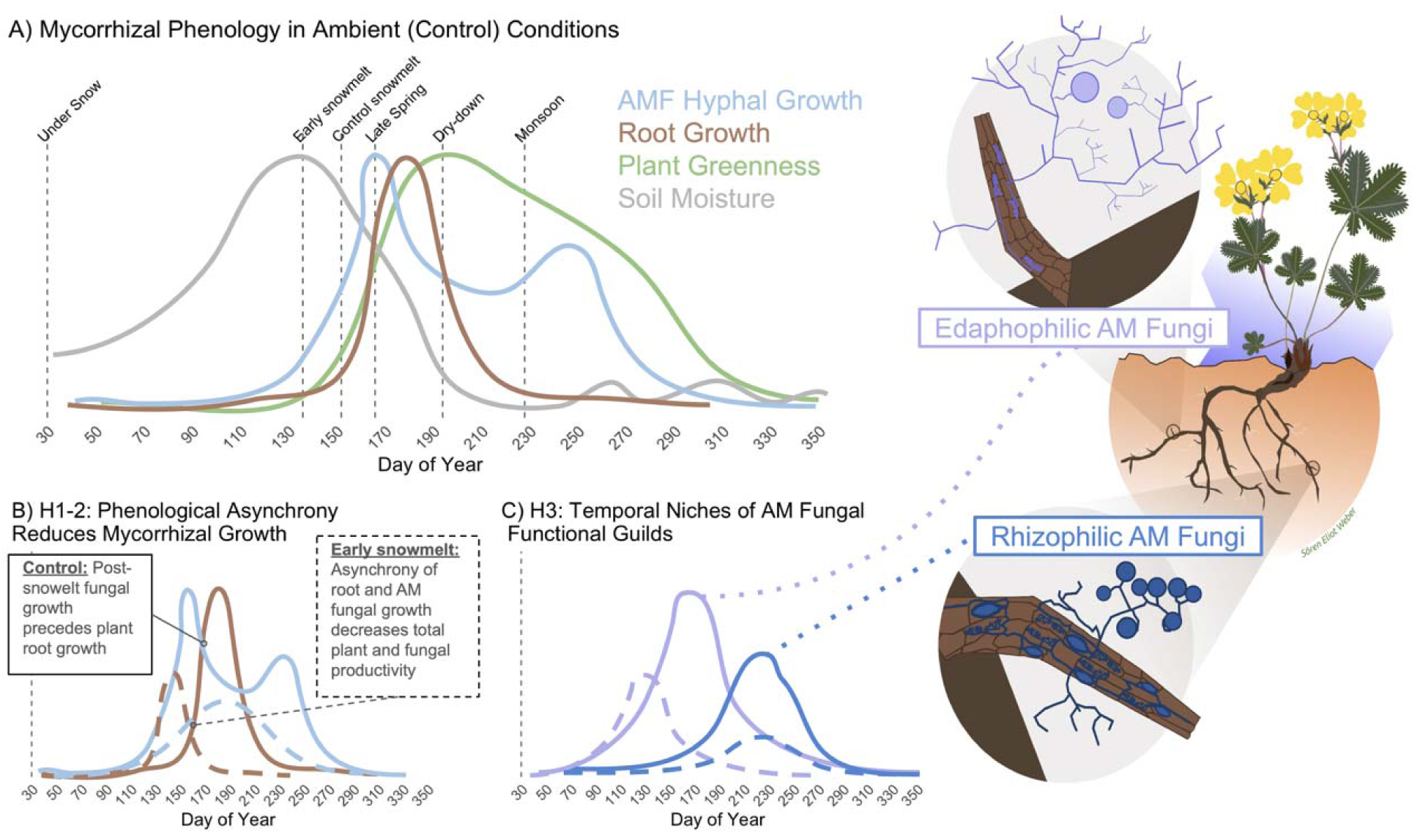
Conceptual Diagram of Mycorrhizal Phenology & Hypotheses. A) Conceptual model of ambient phenological relationships between plant growth, nutrient availability, and soil moisture, based on results in control plots. B) H1-2: Early snowmelt will decouple plant (here shown as root growth) and AM fungal phenology across time, reducing overall mycorrhizal growth. C) H3: Different temporal niches of edaphophilic (soil-colonizing) and rhizophilic (root-colonizing) AM fungi, with different potential responses to early snowmelt.

Warming winters alter snowpack dynamics, reshaping both the timing and magnitude of these microbe-mediated biogeochemical pulses ^12^. When snowpack is reduced or arrives inconsistently, microbial communities lose the protective buffer of the snowpack and experience freeze-thaw cycles, leading to nutrient loss ^13–15^. AM fungi may, alternatively, stabilize nutrient retention and plant–microbe synchrony under changing cold season regimes by supporting multifunctionality under warming and resistance to climatic disturbances ^16^ ^17^. Therefore, AM fungi likely play a central role in buffering alpine ecosystem processes against the destabilizing effects of winter climate change, with their responses to warming likely reshaping N & P cycling in ways that cannot be predicted from plant or climate data alone^18^.

We need to examine the functional traits that may influence AM fungal responses to nutrient availability in order to understand how early snowmelt disrupts plant-AM fungal phenology. Functional diversity of AM fungi may shape community turnover across the growing season, but it is unclear whether their traits follow consistent seasonal niches linked to plant demand or broader ecosystem cycles ^19^, which could impact phenological matching. Trait-based frameworks of AM fungal life history strategies ^20^ may map to phenologies if different AM fungal species are alternatively adapted to tolerating the stress of the cold or drought seasons versus competition for nutrients during the growing season. Traits such as sensitivity to N limitation ^21^ and differences in hyphal allocation strategies ^22^ may further define temporal niches among AM fungi.

Here we apply a trait-based functional guild framework for hyphal allocation strategy to inform community composition and hyphal growth patterns across time. The ERA framework ^22^ classifies AM fungi into three guilds based on hyphal allocation strategy: **E**daphophilic fungi invest heavily in soil-exploring hyphae; **R**hizophilic fungi invest in root colonization; and **A**ncestral fungi exhibit limited hyphal growth in both roots and soil. Because AM fungal guilds differ in hyphal morphology and growth strategies, measuring the diversity of guilds over time provides a sensitive, interpretable metric for detecting shifts in ecological function under climate perturbations^23^. If early snowmelt causes plant-fungal phenological asynchrony, the composition or activity of these functional types may shift in ways that alter ecosystem-scale nutrient cycling, similar to long-term shifts of plant functional types^18,24^. An integrated understanding of how AM fungal traits and nutrient cycling change across seasons is essential for understanding the fate of soil resources under climate change.

*Hypotheses*: In this study, we experimentally advanced snowmelt to test how warming winters influence nutrient cycling through disrupting mycorrhizal processes. We tracked mycorrhizal plant and fungal growth, soil nutrients, and AM fungal community composition to test the following hypotheses: **H1)** Early snowmelt will decouple the growth phenology of plants and their AM fungal partners if they rely on different cues to stimulate growth (Fig 1B). **H2)** Phenological decoupling will reduce plant and AM fungal growth, due to breakdown of plant-fungal nutrient transfer (Fig 1B). **H3)** Early snowmelt will impact ERA Functional guilds to different degrees, because different hyphal growth strategies may have distinct seasonal niches and capacities to align with plant nutrient demand (Fig 1C).

## Materials and Methods

### Field site

We established 5 blocks of 10 x 7 meter plots in a subalpine meadow located within the East River Watershed near Gothic, Colorado (Lat: 38.962396, Long: 106.993848, Elev: 2900m) ^25^. Soil is clay-loam and sandy clay loam mollisols with a plant community dominated by AM fungi-associated perennial herbs, grasses, and shrubs (e.g., *Festuca thurberi*, *Dasiphora fruticosa*). This meadow experiences an average annual precipitation of 88 cm and a growing season spanning from June to September.

### Snowmelt Experiment

The winter of 2022-23 prior to our experiment had a deep snowpack that reached 246 cm, with a total snowfall of 921 cm. Within each block, one plot experienced ambient snowmelt (control plots) and one received an early snowmelt treatment. Snowmelt was advanced in the treatment plots with black shadecloths ^26^. Shadecloths were installed on April 20th when snowpack was 162 cm high, and removed on May 16th when treatment plots were snow-free and control plots had approximately 55 cm of snowpack (Fig 2A). These snowpack measurements were taken at the Gothic weather station (www.gothicwx.org), 45 meters from our plots. The treatment advanced the snowmelt date 12 days ahead of the control snow-free day on May 28th. This advancement simulated 50 years of warming winter based on the observed snowmelt date advancement rate of 2.4 days per decade from 1975-2025 ^27^

**Figure 2:**
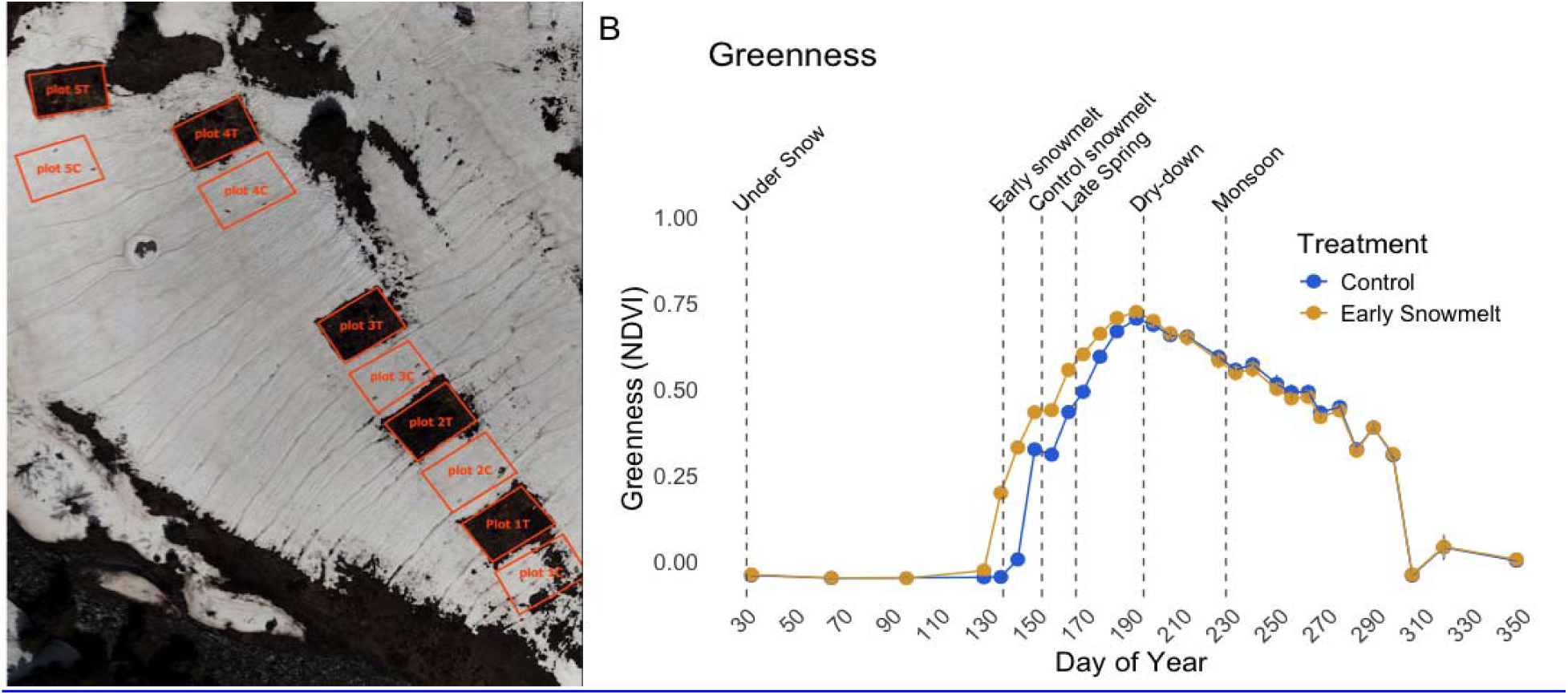
Early snowmelt advances the start of the growing season: A) Plots on May 23, 2023, showing advanced snowmelt in treatment plots. B) Weekly NDVI data shown as mean with standard error. Dotted vertical lines indicate the beginning of each phenological period when soil and ingrowth cores were taken.

### Soil Sampling

We installed ingrowth cores to track root and AM fungal hyphal productivity across the growing season^28^. Soils and nutrients were collected down to 20 cm to capture mycorrhizal dynamics and nutrient fluxes. Soil was collected from the meadow in 2022, sterilized by autoclave, and packed into ingrowth cores with a volume of 364 cm³, holding approximately 320g of soil. 9mm mesh was used to capture root and AM fungal hyphal growth (root ingrowth cores), and 20 micron mesh was used to capture just AM fungal hyphae, excluding roots (root-exclusion ingrowth cores). Three replicates of each core were installed per plot. We placed the first set of ingrowth cores on 9/17/2023, then collected them on the snowmelt date in each plot in the spring of 2023 **(Control: 5/28/23, Treatment: 5/16/23**).

Ingrowth cores were subsequently installed and collected on a monthly basis from May to September. Bulk soil cores (6 x 20 cm) were also collected monthly to measure the standing stock of roots and AM fungal hyphae. Soils were sieved immediately following collection and split into air-dried, 4°C, or -80°C sub-samples for processing. Soils stored at 4°C were processed within 2 weeks of sampling.

To measure dry root biomass, we removed roots from sieved samples for 20 minutes and dried the roots in an oven. To measure AM fungal hyphal soil colonization, we extracted hyphae from 5g of soil with sodium hexametaphosphate, stained hyphal tissue with acid fuchsin, and calculated hyphal length density using the grid line-intercept method ^29^. Root and AM fungal hyphal productivity rates were estimated by normalizing total hyphal length density or root biomass, respectively to the number of days each ingrowth core was in the ground.

### Sensors

TMS-4 sensors (TOMST, Praha, Czech Republic) were installed in each plot to take continuous measurements of soil moisture (% volumetric soil water content, VWC) and temperature (°C) at 6cm deep, as well as air temperature above soil surface at 2 and 15 cm (sFig 1). Probes were installed in the summer of 2022, with readings taken every 15 minutes throughout the growing season. To detect above-ground plant growth phenology (Fig 2B), we measured greenness by extracting Normalized Difference Vegetation Index (NDVI) values from weekly imagery collected of all plots with a Micasense Altum V1 multispectral and thermal sensor mounted on an uncrewed aircraft system.

### Soil Nutrients

Soil pH was measured on a slurry of 10 g dry soil in 50 mL deionized water using a calibrated pH meter. Percent of soil organic matter (SOM) was measured using the loss-on-ignition method ^30^, at 360°C for 2 hours. We extracted 5 g of fresh soil with K□SO□ to quantify the total pool size of soil-extractable NH_4_^+^ and NO_3_^-^. The soil extracts were filtered and then stored frozen until analysis. Soil nitrate concentrations were measured using the vanadium chloride colorimetric method ^31^, and ammonium concentrations were determined using the indophenol blue colorimetric method ^32^. Dissolved organic carbon was quantified using the Mn(III)-pyrophosphate colorimetric assay ^33^.

### Ion Exchange Resins

We incubated and periodically collected ion exchange resin capsules to estimate net soil nutrient fluxes. In each plot, we installed two custom-built WECSA-type soil access tubes ^34^. The soil access tubes were made using a 1.5″ (∼4 cm) diameter PVC outer sleeve and a removable 1″ (∼2.5 cm) diameter PVC inner sleeve inserted at a 45° angle to 20 cm depth below the surface. A Unibest PST-1 resin capsule (Unibest International) was securely attached to the inner sleeve and kept in direct soil contact using a 1 to 0.75” PVC reducer bushing that was drilled with ⅝” holes to allow soil pore water flow through. Resin capsules were installed and collected in time with ingrowth core collections (Fig 4) to measure available nutrients. Net nutrients fluxes were estimated by extracting ion exchange resin capsules with 25 mL of 1M HCl. The resin extracts were vacuum-filtered across 0.2 □m membranes using Acroprep 24-well filter plates (Cytiva). The concentration of NH_4_^+^ and NO_3_^-^ in the resin extracts was determined colorimetrically as described for soil extracts. Resin-extractable phosphorus was determined using an Agilent 8900 Inductively Coupled Plasma Mass Spectrometry (ICP-MS) by diluting resin extracts 1:100 (vol/vol) in 2% nitric acid and quantifying total P (parts per billion). The estimated net nutrient flux (nmol day^-1^) was calculated by normalizing the nutrient concentrations and dividing by the number of incubation days in the field.

### AM Fungal sequencing and bioinformatics

DNA was extracted from ∼0.2g of frozen soil using the Qiagen Dneasy Powersoil Pro kit. AM fungi were amplified with NS31/AML2 primers ^35^ and sequenced on the Illumina NovaSeq pair-ended 2 X 300 bb reads. Amplicon sequence variants (ASVs) were classified using QIIME2 and the MaarjAM database ^36,37^. ASVs were merged at the MaarjAM-VTX level to analyze communities over time. AM fungal reads were also classified into the ERA (edaphophilic, rhizophilic, or ancestral) trait-based framework for hyphal allocation strategy based on family-level taxonomy^22^.

### Statistical analysis

All statistical analyses were performed in R (v4.5.1). We determined AM fungal diversity with calculations of q0 species richness and q1 Shannon’s exponent. We used the nlme package to run mixed effects linear models (MELMs) with random block effects nested within a 1st order temporal autocorrelation structure (TAC) to determine the effect of early snowmelt on response variables over time, with post-hoc tests for treatment at each timepoint (sTable 1). We also used an MELM with TAC to determine the edaphic drivers of root productivity, AM fungal hyphal productivity, and NDVI (sTable 2). For this MELM with multiple effects, model selection was performed with stepwise AIC. For all MELM models, co-linearity was assessed with the vif function. We determined effect sizes using standardized beta coefficients (□) calculated with the standardize_parameters function.

To analyze the AM fungal community, we performed distance-based redundancy analyses on the effects of time and treatment (sTable 3) and the effects of edaphic factors on functional guild composition (sTable 4). AM fungal abundances were center-log normalized and used to calculate Aitchison distance matrices ^38^. We ran dbRDA models to test effects on AM fungal composition, using stepwise AIC regression to find the best-fitting models.

## Results

### 1. Advancement of snowmelt date and plant phenology

#### Review of ecosystem phenology

Based on soil moisture trends, greenness trends, and the intervals of belowground growth we captured with ingrowth cores, periods of the growing season will be referred to as: 1) **Under-snow**: 09/17/23 to Snowmelt; 2) **Post-Snowmelt**: Snowmelt-06/15/23; 3) **Late Spring**: 06/15/23-07/13/23; 4) **Dry-down**: 07/13/23-08/16/23; 5) **Monsoon:** 08/16/23-09/09/23 (Fig 2B).

Greenness rapidly increased post-snowmelt until precipitation ceased at the beginning of the dry-down period, peaking ∼40 days after snowmelt, then steadily decreasing through the dry-down and monsoon periods (Fig 2B). Root productivity and standing stock peaked before greenness, in the late spring period (Fig 3E-F). AM fungal productivity peaked before roots in the post-snowmelt period, in the 14-28 days after snowmelt (Fig 3B). Standing stock of AM fungal hyphae peaked at the beginning of the season (Fig 3C), indicating a high rate of turnover following snowmelt. We observed substantial AM fungal hyphal density under snow, (1-3 m hyphae / g soil, sFig 2A) even though the estimated productivity was lower.

**Figure 3:**
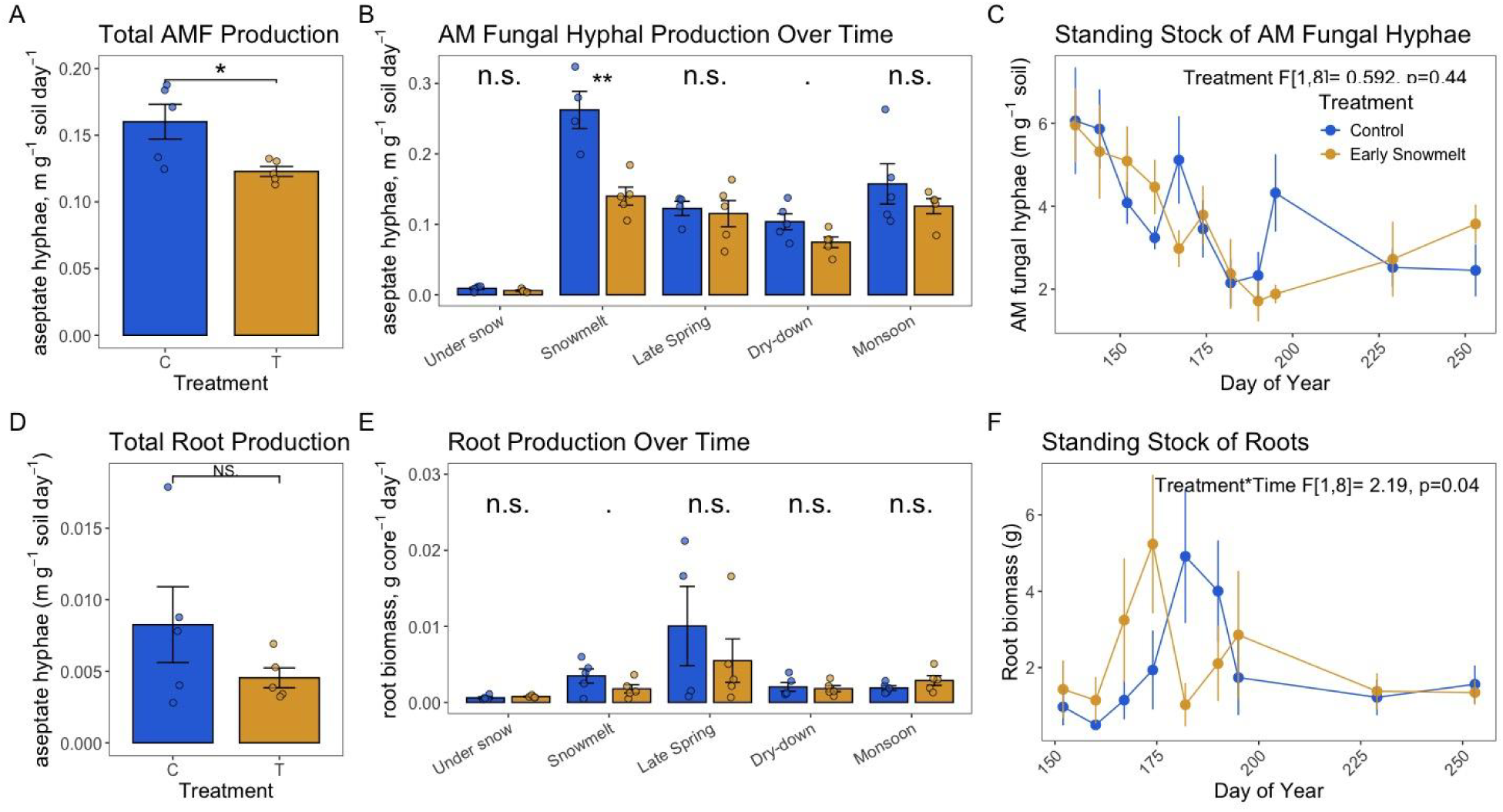
Early snowmelt reduces AM fungal production: A,D) Cumulative productivity averaged over the entire season. B,E) Estimated productivity for each growing period. Mean with standard errors is plotted as a bar, with post-hoc significance testing. Stars indicate significance level (. = <0.10, * = <.05, ** = <.01, *** =<.001). C,F) Standing stock over time, with early snowmelt treatment and time effect from MELM.

### Early snowmelt results

Advancing snowmelt by 12 days (Fig 2A) strongly affected soil moisture (F[1,4]=8.357, p=0.009), increasing soil moisture by 2-3% before snowmelt and decreasing soil moisture post-snowmelt, including late into the growing season during the monsoon period (sFig 1). Early snowmelt lead to earlier warming of air and soil temperatures, starting after the snowmelt date (sFig 1). Early snowmelt treatment advanced onset of greening by ∼10 days, with treatment plots significantly greener earlier in the growing season compared to control (Fig. 2B, sTable 1, under-snow p= 1.60E-05, post-snowmelt p=0.002). Plant greenness trends in all plots converged in mid July, coinciding with onset of the dry-down period before the arrival of the monsoon.

### 2. Root & Hyphal Responses to early snowmelt

Early snowmelt decreased the average rate of AM fungal hyphal production across the whole season by 23% (Fig 3A, F[1,8] = 7.53, p=0.02, □=-1.32). AM fungal hyphae grew 46% less immediately post-snowmelt and 28% less during dry-down (Fig 3B, sTable 1). The hyphal density inside ingrowth cores taken on the snowmelt date was also 38% lower in treatment plots, by 0.5 m per g soil (F[1,4] = 5.233, p=0.08, sFig 2A). We observed similar trends with weaker effects in the root exclusion cores (sFig 2B). Early snowmelt did not affect the AM fungal hyphal standing stock (Fig 3C), suggesting that reduced production may have been offset by changes in turnover.

Root growth was on average 44% lower in early snowmelt treatment plots, but these differences were not statistically significant (F[1,8] = 1.84, p = 0.21, □ = -0.82; Fig. 3D-E). Treatment and time interacted to advance peak root standing stock by ∼1 week in early snowmelt plots (Fig. 3F), suggesting that snowmelt cues on plant growth are symmetrical above- and belowground.

In summary, early snowmelt decoupled plant and AM fungal growth. Early snowmelt advanced plant greenness (Fig 2B) and root standing stocks (Fig 3F) while decreasing AM fungal hyphal productivity (Fig 3A-B). Effects were observed in the post-snowmelt period in early June, indicating that the liminal period of mycorrhizal growth when production shifts from extraradical hyphae to roots is especially vulnerable to warming winter.

### 3. Edaphic responses to early snowmelt and their relationships to mycorrhizal growth

Early snowmelt affected soil nutrients during the post-snowmelt window, coinciding with less AM fungal hyphal growth (Fig. 4, sTable 1). Early snowmelt treatment during this early window resulted in 21% less SOM (Fig 4F, □=-1.66), 30% less available NH_4_^+^ (Fig 4A, □=-1.41), 22% less available P (Fig 4C, □=-1.30) and 11% less total soil NO_3_^-^ (sFig 3B, □=-1.59). Early snowmelt increased total NO□□ from undetectable in controls to 0.154 nmol g□¹ soil (sFig 3, F[1,5] = 3.518, p = 0.034, □ = 1.05) and total NH□□ by over sixfold (sFig. 3, F[1,5] = 13.152, p = 0.022, □ = 1.49), demonstrating accelerated N mineralization under snow. Soil pH varied over time (5.6-6.9, sFig 3C) but was not influenced by early snowmelt treatment (sTable 1).

**Figure 4:**
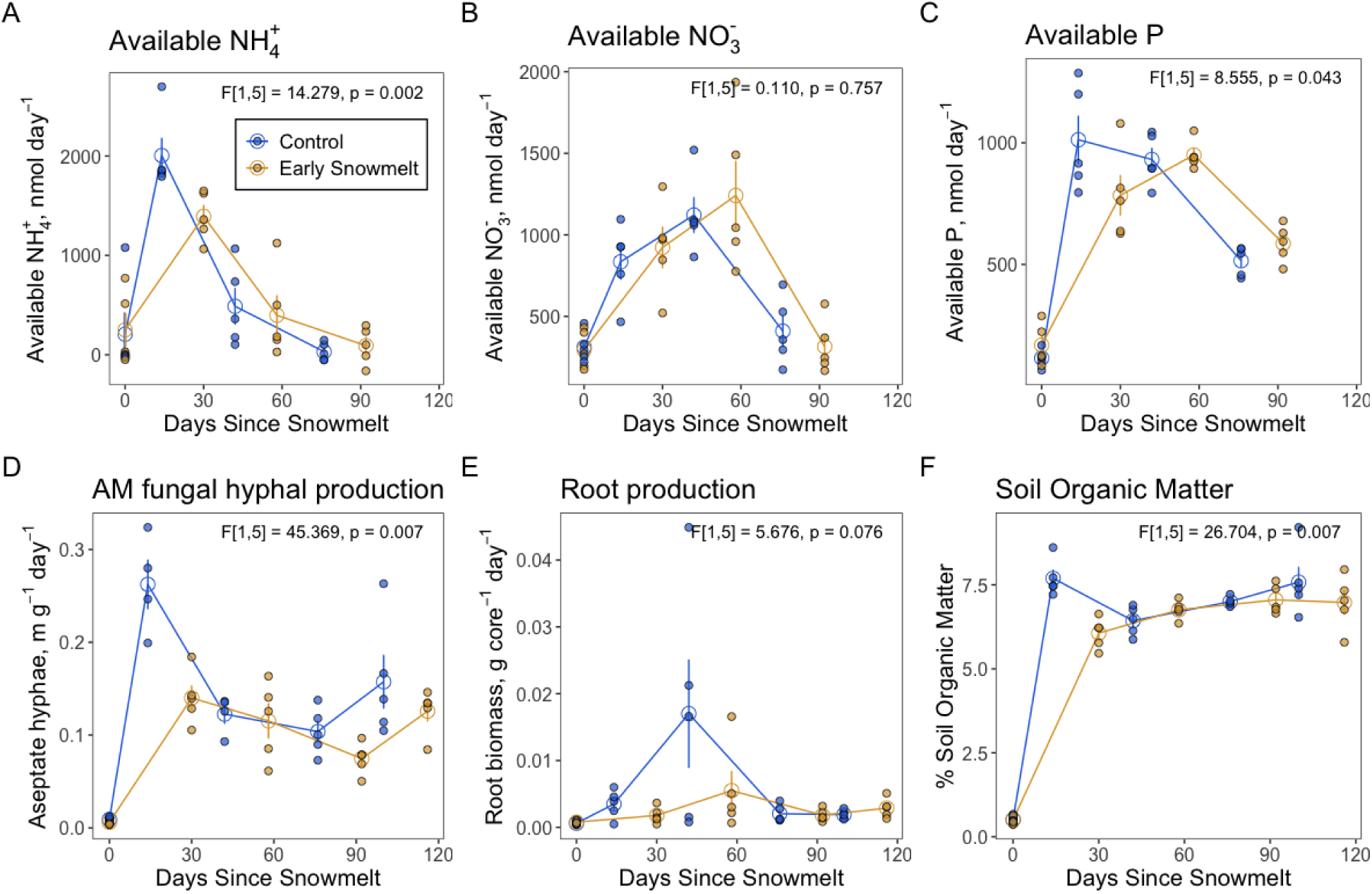
Early snowmelt changes nutrient cycling: Concentrations of available nutrients measured with resin exchange capsules (A-C), growth rates from belowground growth ingrowth cores (D-E), and % soil organic matter in ingrowth core (F). Treatment effects for the post-snowmelt period are shown on each plot, full models are in sTable 1.

The AM fungal hyphal growth rate was positively correlated with available NH_4_^+^ (F[1,6] = 55.541, p<.0001, □ = 0.95) and P (Fig 4A, (F[1,6] =4.397, p=0.081, □ = 0.40), linking lower AM fungal hyphal growth to lower nutrient concentrations post-snowmelt (sTable 2). Early snowmelt potentially weakened the coordination of root–fungal nutrient exchange, as there was 8.4% less available P per meter of hyphae and 27.1% less hyphae per unit root biomass (sFig 4), suggesting disrupted phosphorus transfer between roots and AM fungi. The strong negative correlation between greenness and available NH□□ (F[1,10] = 69.23, p < 0.0001, β = –1.13; sTable 2) suggests that plant growth results in depletion of nutrients from the soil pool across time.

### 4. Functional traits of AMF

Overall, AM fungal guilds with different hyphal allocation strategies displayed distinct temporal patterns of diversity, composition, and responses to early snowmelt (Fig 5, sTable 3). We found that Edaphophilic AM fungi dominated in the early season, while Rhizophilic AM fungi diversified later in the season, after root development. Ancestral AM fungi were rare (<30 ASVs). These fungi may be most active during winter, consistent with previous findings that ancestral AM fungi dominate at higher elevations where soils are colder, wetter, and more acidic ^39^.

**Figure 5:**
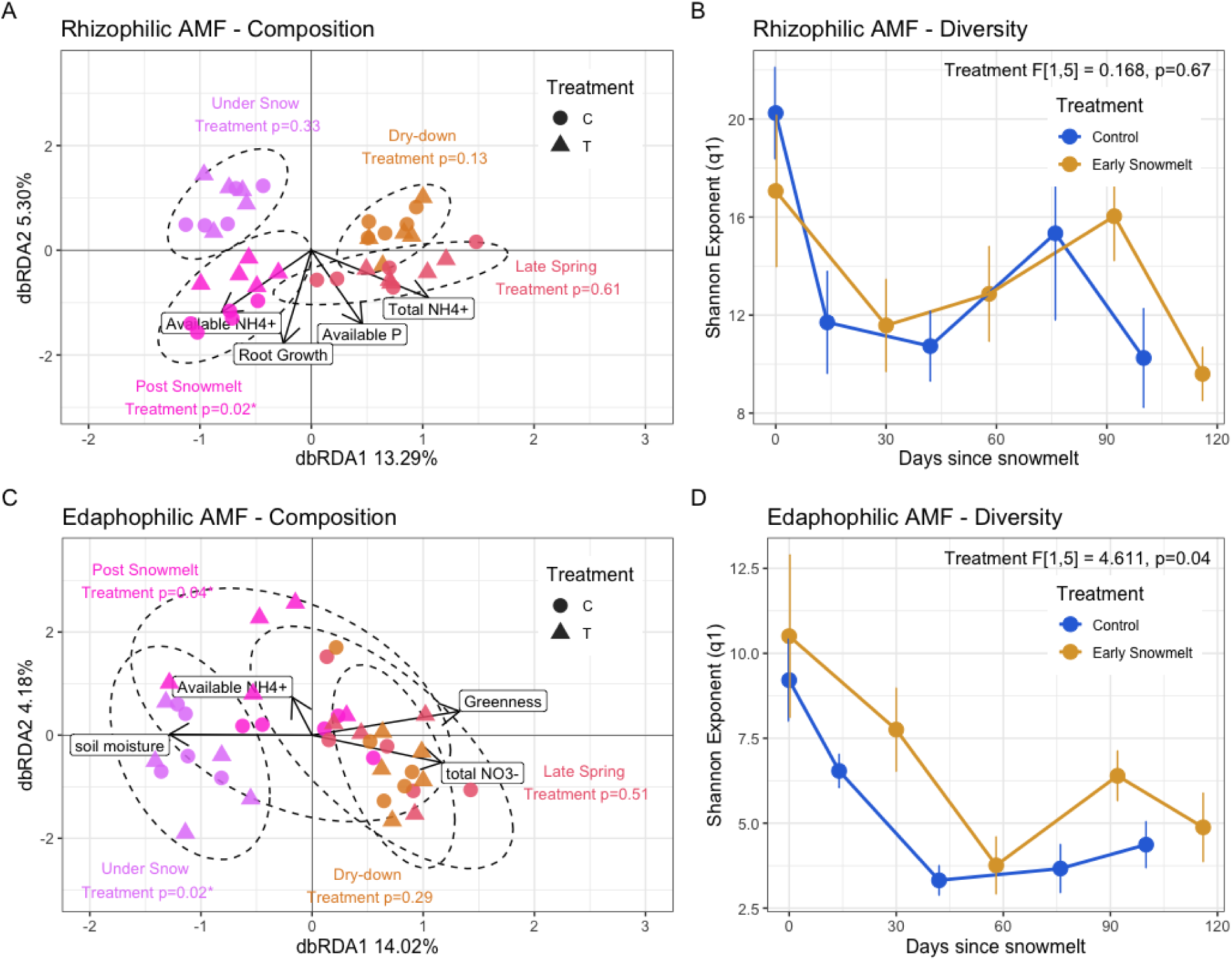
Hyphal allocation strategy influences AM fungal phenology and response to early snowmelt. Composition dynamics shown for both Rhizophilic and Edaphophilic functional guilds of AM fungi. Constrained ordinations were made with arrows representing significant drivers of composition from dbRDA (A,C). Warm colors and ellipses represent time periods across the growing season, and ellipses are annotated with treatment effects. Diversity over time is also shown for each functional group (B,D) and annotated with treatment effect from MELM. Error bars show standard error.

The root-colonizing rhizophilic group peaked in diversity during the dry down period (Fig 5B), indicating diversification after roots grow and uptake nutrients. Temporal turnover of rhizophiles (Fig 5A) was driven by root growth (F[1,29]=1.829, p=0.018) and available NH_4_^+^ (F[1,29]=3.104, p=0.001). Early snowmelt shifted rhizophilic composition after snowmelt (F[1,38] = 4.42, p = 0.018), coinciding with lower NH□□ and root biomass (Fig. 5B), suggesting root growth limitation influenced rhizophilic fungi.

The soil-colonizing edaphophilic group peaked in diversity during snowmelt, steadily declining as the season progressed (Fig 5D). Temporal turnover of edaphophiles (Fig 5C) was driven by greenness (F[1,29]=2.541, p=0.036) and increasing levels of total soil N (F[1,29]=1.826, p=0.024). Early snowmelt increased edaphophilic diversity and altered community composition under snow and post-snowmelt (Fig. 5C–D, sTable 1). This pattern may reflect greater resource allocation to nutrient-foraging edaphophiles due to faster depletion of nutrients by early plant growth, but hyphal biomass did not increase (Fig. 3A–B), indicating that edaphophilic shifts did not offset overall declines in AM fungal growth.

## Discussion

*Overview of findings:* We found that early snowmelt advanced the onset of plant greening (Fig 2) and peak root standing stock (Fig 3), but reduced AM fungal hyphal production (Fig 3, 4).

Early snowmelt reduced soil organic matter, P, and NH_4_^+^ post-snowmelt (Fig 4), suggesting that early-greening plants may have quickly depleted nutrient pools. The AM fungal community changed with early snowmelt (Fig 5), indicating that differences in total hyphal growth was mediated by shifts in AM fungal species and diversity. Furthermore, functional guild analysis revealed that AM fungal species had unique temporal niches cued by the timing of nutrients, root growth, and greening (Fig 5). Overall, we found support for our hypotheses (Fig 1), and strong evidence that early snowmelt decouples plant and AM fungal growth at the ecosystem scale. Nutrient pools and functional guild analysis of the AM fungal community provide further clues to the mechanism underlying phenological asynchrony.

### Early snowmelt temporally decouples plants from mycorrhizal fungal partners

Lower AM fungal growth could be caused by less resource allocation to mycorrhizal partners, if the shorter photoperiod and colder temperatures experienced by early greening plants constrained photosynthesis and increased frost exposure, limiting host resource allocation ^40–42^. However, during this experiment we also found that early snowmelt increased plant productivity ^43^, indicating that earlier growth onset doesn’t decrease, but *increases* carbon fixed by the host plant. Our results show that faster early-season plant growth coincides with reduced AM fungal growth, indicating a mismatch between host plant demand and fungal nutrient supply. These findings support our hypothesis that early snowmelt decouples the phenology of plants and their AM fungal partners (**H1**).

Decoupled plant-mycorrhizal fungal phenology may parallel mechanisms of plant asynchrony with pollinators, herbivores, and even fungal pathogens. For example, early snowmelt has been shown to advance plant phenology, but not pollinator insect emergence, due lagged soil warming ^44^. Mycorrhizal phenology may also share similarities in temporal dynamics with fungal pathogens, where infection of plant tissues depends on seasonal patterns of plant resource allocation to growth versus defense^45^. Our observations suggest that like other plant symbioses, different environmental cues and life-history trade-offs create baseline asynchrony between plant and fungi that may make symbiotic relationships vulnerable to climate change^46,47^. Early snowmelt effects were strongest early in the season, with plants and fungi showing weak or no treatment response by fall. Over longer timescales, warming winters can increase AM fungal hyphal production and restore temporarily depleted soil C stocks ^24,48^ Therefore, short-term asynchrony may eventually stabilize, but with high potential for restructured, non-analogous communities shaped by new traits and phenological dynamics^49^.

### Phenological asynchrony disrupts plant-fungal nutrient cycling, reducing mycorrhizal productivity

We found that early snowmelt reduced AM fungal hyphal production and available N and P, supporting our hypothesis that phenological decoupling would decrease mycorrhizal growth through disrupted nutrient flow **(H2)**. Early snowmelt caused belowground root development to advance with aboveground greening, but decoupled and decreased AM fungal hyphal production. The causal relationship between lower soil nutrients and less AM hyphal production is unclear, likely reflecting multiple mechanisms of mycorrhizal response to warming winter.

Early snowmelt may have reshaped the soil microenvironment in ways that suppressed hyphal growth and foraging. Under snowpack, microbes decompose organic matter and mineralize nutrients ^9,50^, providing N and P that AM fungi access under snow. Early snowmelt reduces microbial biomass and increases denitrification, increasing nutrient loss ^13^. If early snowmelt shrank the nutrient pool size, this may have weakened the mycorrhizal symbiosis, because insufficient nutrient supply from fungus to plant can trigger arbuscule degradation and stagnate hyphal networks ^51^. Weakened AM fungal hyphal networks during the post-snowmelt period could further reduce nutrient retention and stability by disrupting hyphosphere interactions and nutrient□retaining microsites, causing nutrient pulses to become mistimed with mycorrhizal uptake ^13,52^.

However, if early snowmelt causes nutrient limitation via nutrient loss, stronger plant-fungal coupling might be expected based on functional equilibrium models for mycorrhizal function ^53,54^. Instead, advancing plant growth may have resulted in direct uptake of nutrients with limited hyphal development. Early greening plants may have aligned their nutrient demand more closely with labile nutrient mineralization from overwintered microbial biomass and other detritus^55^. Under ambient conditions, AM fungi may bridge the temporal gap between nutrient release under snowpack and root demand post-snowmelt ^8^. If early snowmelt compresses these phenological windows, plant-fungal complementarity could be weakened ^56^. In montane ecosystems with short growing seasons, temporal complementarity between plant hosts and mycorrhizal fungi is probably critical to maintain resource use efficiency, as plants benefit from symbionts that extend nutrient uptake during late-season cold or early-season snow cover.

### Hyphal allocation strategy shapes temporal niches and responses to early snowmelt

We used a trait-based guild framework to determine how hyphal allocation strategy (Fig 1) influences AM fungal responses to early snowmelt (Fig 5). We found that edaphophilic and rhizophilic AM fungi had different responses to early snowmelt, supporting our hypothesis (**H3**) and confirming other findings of taxa-specific responses to warming ^57^. The soil-colonizing edaphophilic AM fungi decreased in diversity and shifted in composition post-snowmelt, indicating that treatment effects on nutrient-foraging mycorrhizal function is related to plant-fungal asynchrony. These findings support other evidence that soil-foraging edaphophilic AM fungi are often more vulnerable to shifts in nutrient dynamics compared to rhizophilic types ^21,58,59^. Root colonization was not measured in this study, which might have revealed more rhizophilic responses to early snowmelt. Some AM fungal groups may be especially vulnerable to warming winters, and these results demonstrate how trait-based understanding of mycorrhizal communities can enhance predictions of ecosystem responses to warming. The ERA framework ^22^ is a taxonomy-based way to group AM fungi, but taxonomic identity alone does not consistently predict functional responses across environmental gradients ^60,61^. A suite of traits including hyphal length and branching, sporulation, nutrient transporter expression, and host specificity define life-history strategies that drive responses to warming ^20,62^. These life-history strategies may ultimately govern not just mycorrhizal fungal persistence and phenology across the growing season, but the ability of the symbiosis to buffer nutrient losses and sustain plant communities under changing winter conditions.

Also supporting our third hypothesis, diversity analysis showed that AM fungal guilds exhibited distinct temporal niches across the growing season (Fig 5). Edaphophilic fungi dominated early when hyphal productivity (and likely, host nutrient demand) was high. Rhizophilic fungi diversified later after peak root growth. These findings indicate that host plant phenology, environmental conditions, and fungal traits together partition mycorrhizal niches across time.

This species turnover across the growing season also indicates that the AM fungal community is more temporally dynamic than is often assumed ^63^. If this trend is applicable to other systems, single timepoint studies may underestimate the effects of global change on AM fungal composition and function. Seasonal environmental changes likely alter plant–fungal associations in ways similar to climate-driven shifts in nutrient availability ^64,65^. Notably, we discovered that when the greening rate is highest in the middle of the season (Fig 1), the AM fungal community reaches the lowest species diversity (sFig 5) and no treatment effects are observed on any responses. We observed highest AM fungal diversity and treatment responses in the early season. Therefore, the effects of climate change on the AM fungal symbiosis may be strongest outside of peak plant growth, in the leading and lagging periods where mycorrhiza perform key belowground functions in response to host phenophases.

## Conclusions

By experimentally advancing snowmelt in a sub-alpine meadow, we demonstrate that warming winters cause phenological asynchrony between host plant and arbuscular mycorrhizal fungi. Early snowmelt caused early onset of greening and diminished extraradical hyphal production by mycorrhiza. Plant-fungal asynchrony was accompanied by smaller pools of available P and N, suggesting early, direct nutrient adsorption by plants lowered host investment in mycorrhizal fungal partners. The mycorrhizal symbiosis was weakened during a critical period of nutrient exchange. These findings point to the vulnerability of belowground mutualisms to warming winters, and highlight the need to incorporate fungal phenology into predictions of ecosystem responses to climate change. Future work scaling early snowmelt effects on belowground microbes to watershed N & P dynamics, or loss of N to the atmosphere, may provide a clearer picture on the relationships between warming winter, nutrient cycling, and plant-fungal asynchrony. The AM fungal community was highly variable across the growing season, indicating that mycorrhizal responses to warming are context-dependent on seasonal variation of soil resources and trait-based community assembly. Together, our results show that warming winters disrupt the seasonal timing of plant–fungal interactions, potentially triggering nutrient-limitation feedbacks with potential consequences that extend from soil microbial processes to watershed-scale N and P cycling. By revealing both the sensitivity of belowground mutualisms to warming and the strong seasonal shifts in AM fungal community structure, this study underscores the importance of integrating fungal phenology, microbial functional traits, and nutrient-cycling mechanisms into predictions of ecosystem stability under continued climate change.

## Acknowledgements

This research was funded by the U.S. Department of Energy, Office of Science, Office of Biological and Environmental Research, Terrestrial Ecosystem Sciences program under award number DE-FOA-0002392.

We thank billy barr for providing the weather station data.

We thank Soren E. Weber for providing thoughtful feedback on the manuscript, and creating the included illustrations for the ERA framework functional guilds.

## Competing Interests

The authors declare that they do not have any conflict of interest.

## Author contributions

SK, HS, AC, PS, and DI designed the study. HS, PF, OV and PS performed the fieldwork and processed soils and roots in the laboratory. IB and AH operated the drone and analyzed the drone data. HS, PS, ES, and WD performed soil chemistry analyses. HS analyzed the data and wrote the manuscript. All authors contributed to data interpretation and approved the manuscript.

## Data Availability Statement

The data that support the findings of this study are openly available in the Environmental System Science Data Infrastructure for a Virtual Ecosystem (ESS-DIVE). Data can be found on the ESS-DIVE archive (doi:10.15485/2568071). AMF fungal SSU sequences were archived at [NCBI SRA submission in process, database ascension ID will be provided upon manuscript acceptance].

## Notes

### Competing Interest Statement

The authors have declared no competing interest.

