## Supplemental Figures for "Warming winter disrupts mycorrhizal phenology and plant-fungal nutrient cycling"

1. Soil moisture & temperature: Grey line = tarps deployed onto treatment plots. Yellow line = snow-free day of treatment plots, Blue line = snow-free day of control plots. Circular points show mean across plots and plus signs show individual data points.

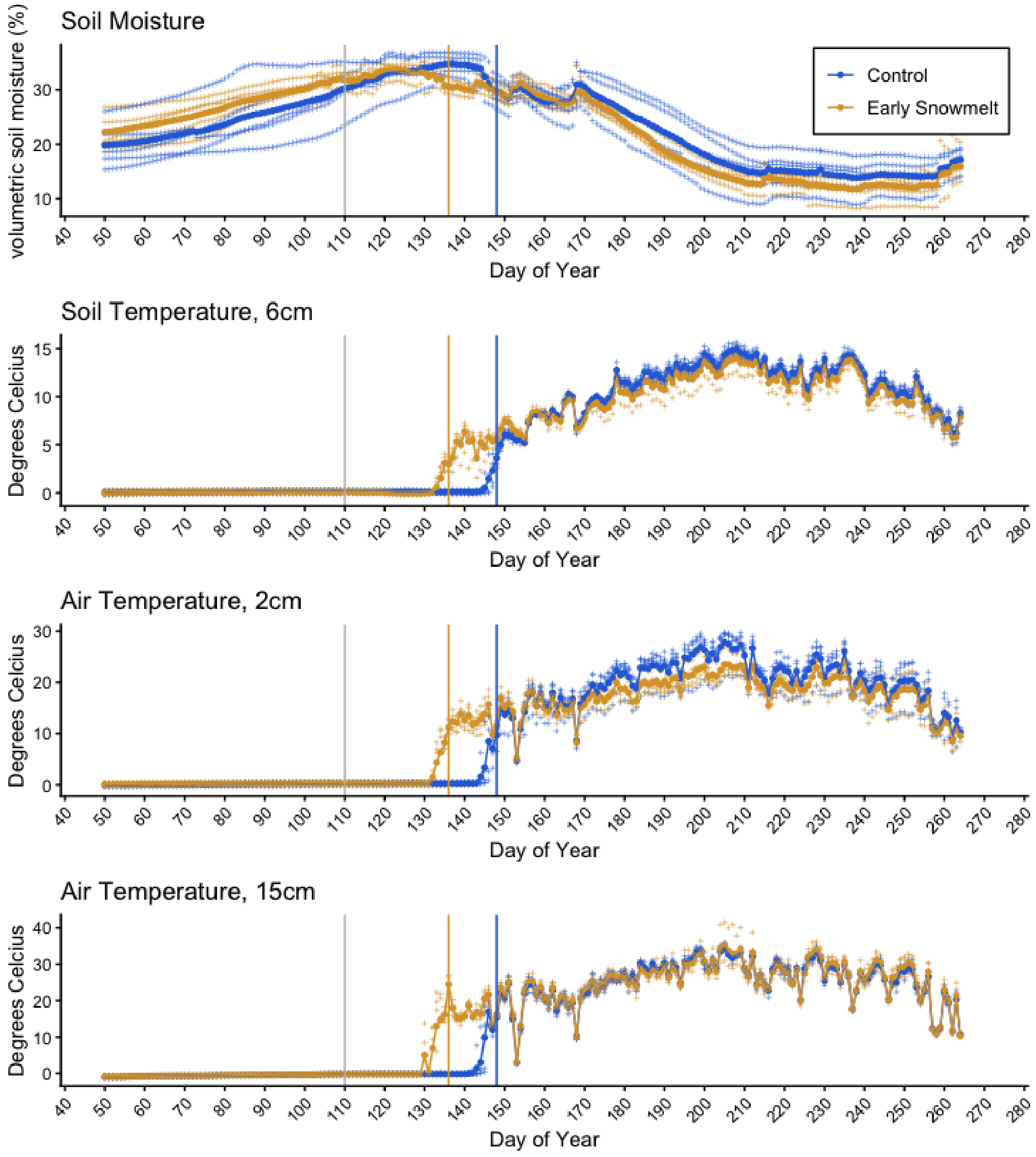

2. Ingrowth core data: Hyphal density and root biomass in each ingrowth core at each sampling period. Note that the snowmelt date for the control (blue) and treatment (yellow) groups is different by 12 days, which is why data is normalized to length of growing period in the main text figures. Mean with standard error is plotted, with individual data points overlaid.

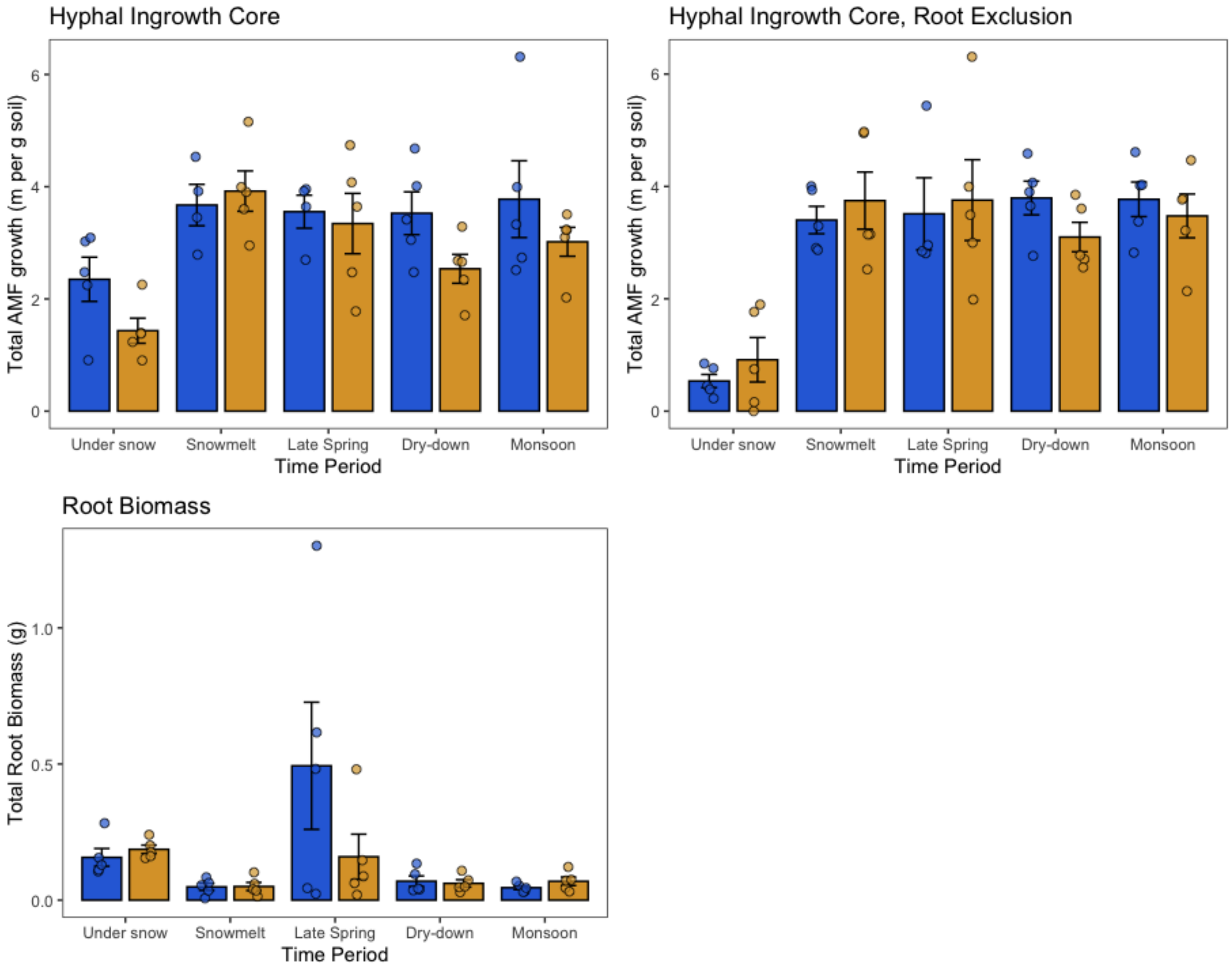

3. Supplemental Soil Chemistry Data: Total soil-extractable nitrogen is shown along with pH. These nitrogen measurements were taken from K2SO4 extract of the entire root ingrowth core, main text figures show available N data from resin exchange cores. Mean with standard error is plotted with separate lines for control (blue) and treatment (yellow).

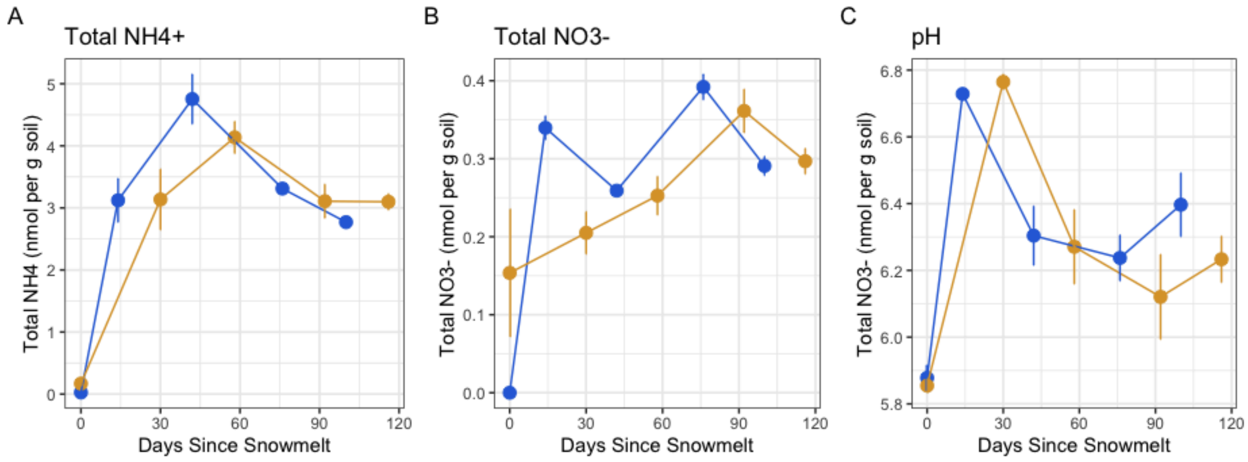

<u>4. Early snowmelt changes mycorrhizal phosphorus cycling and resource allocation.</u> Linear relationships are shown between AM fungal hyphae and available P (A) and root biomass (B). MELM with temporal autocorrelation analysis (sTable 2) revealed that these relationships were significantly influenced by treatment.

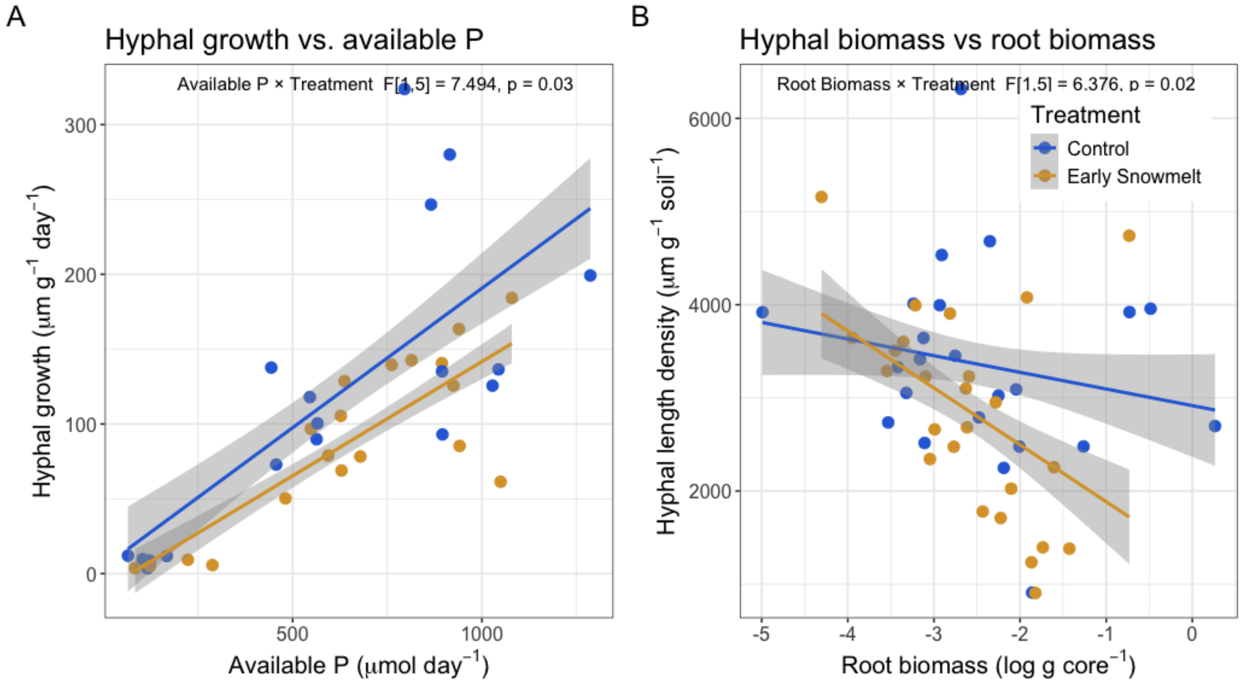

5. Proportional abundance of individual AMF genera. Mean and standard error plotted for the cumulative abundance of each genus across time.

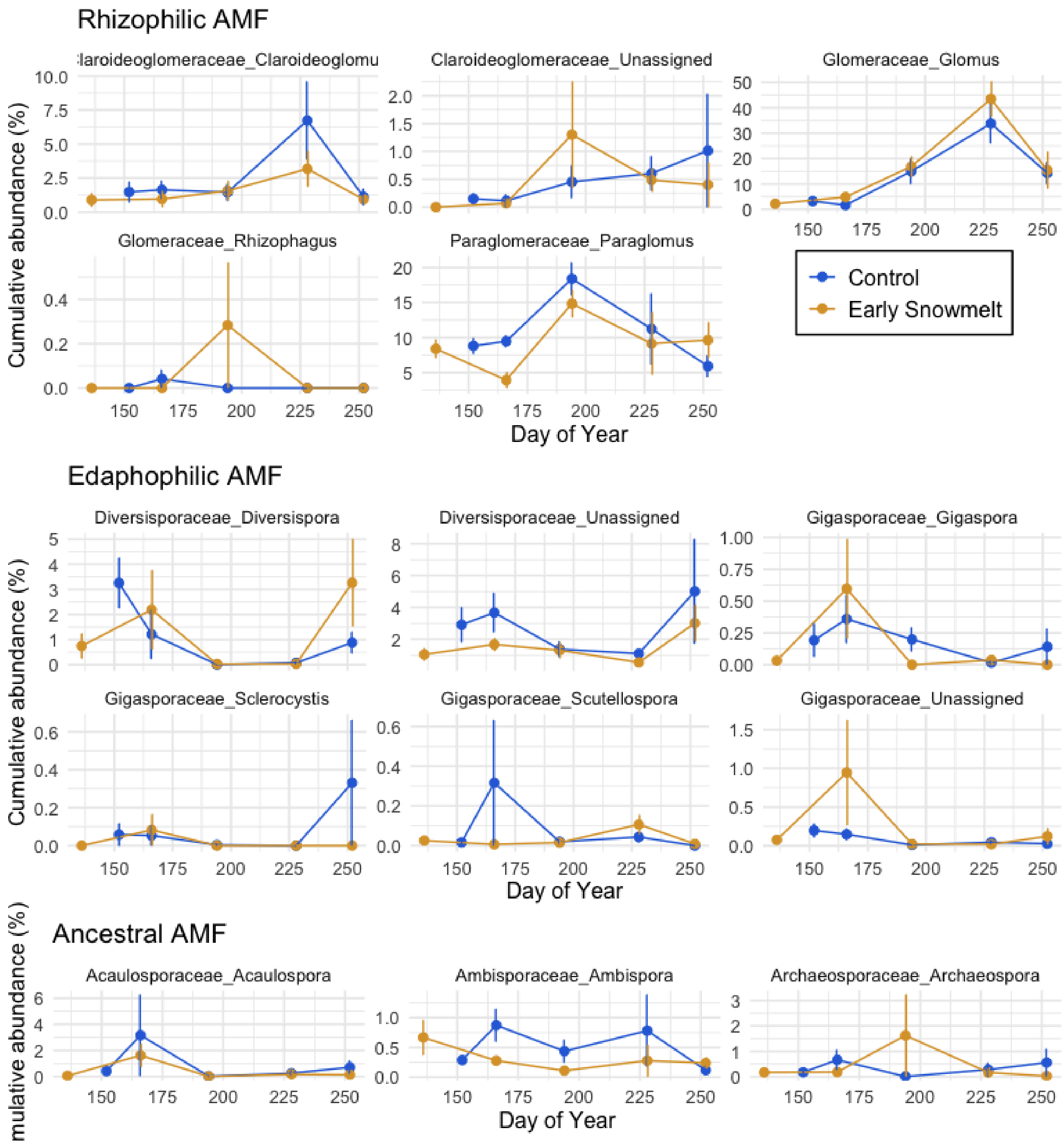
