## Supplemental Tables for "Warming winter disrupts mycorrhizal phenology and plant-fungal nutrient cycling"

| | |
|---|---|
| Table 1 | Mixed effects linear model for effects of time and early snowmelt treatment on responses. F= F-value, p = p-value, B = effect size measured as standardized beta value, Df = degrees of freedom, Int = intercept. Post-hoc tests on treatment effect ran for each sampling period. Model form = response ~ Week + Treatment, random = 1\|Block / Week |
| Table 2 | Mixed effects linear model with temporal auto-correlation structure to determine treatment effects on edaphic drivers of plant and hyphal growth. F= F-value of effect, p = p-value of effect, B = effect size measured as standardized beta value, Df = degrees of freedom, Int = intercept. Model form = response ~ (effects) + 1\| Block + TAC. Stepwise regression used to select effects based on biological hypotheses. |
| Table 3 | Distance-based redundancy analysis for effects of time and early snowmelt treatment on AMF composition. Results are shown for total community, the 3 ERA guilds, and the remaining AMF ASVs which could not be assigned family-level taxonomy. Model form = feature table ~ Condition(Block) + Treatment |
| Table 4 | Distance-based redundancy analysis to determine edaphic drivers of AMF community composition. Model form = feature table ~ Condition(Block) + (effects). Stepwise regression used to select effects based on biological hypotheses |

| | | MELM Results | | | | | | | Post-hoc tests | | | | | | | | | | | | | | |
|---|---|---|---|---|---|---|---|---|---|---|---|---|---|---|---|---|---|---|---|---|---|---|---|
| | | Treatment | | Time | | | | | Under Snow | | | Post-Snowmelt | | | Late Spring | | | Dry Down | | | Monsoon | | |
| | ~ | F | p | F | p | Int | Df | R2 | F | p | *B* | F | p | *B* | F | p | *B* | F | p | *B* | F | p | *B* |
| Growth | AMF productivity | 14.125 | 0.002 | 25.361 | 5.12E-08 | 388.15 | 5 | 0.750 | 4.174 | 0.111 | -1.02 | 45.369 | 0.007 | -1.71 | 0.102 | 0.771 | -0.23 | 4.641 | 0.098 | -1.15 | 1.080 | 0.357 | -0.65 |
| | Root productivity | 2.387 | 0.139 | 3.544 | 0.019 | 17.57 | 5 | 0.291 | 1.273 | 0.322 | 0.70 | 5.676 | 0.076 | -0.92 | 1.815 | 0.249 | -0.82 | 0.140 | 0.727 | -0.22 | 3.650 | 0.129 | 0.83 |
| | Greenness (NDVI) | 4.355 | 0.051 | 115.793 | 4.66E-14 | 5094.82 | 4 | 0.920 | 608.869 | 1.60E-05 | -1.84 | 53.514 | 0.002 | 1.77 | 0.909 | 0.394 | 0.61 | 0.116 | 0.751 | -0.23 | 0.213 | 0.669 | -0.31 |
| Edaphic Measurements | Soil Moisture (VWC) | 8.357 | 0.009 | 105.611 | 1.13E-13 | 795.55 | 4 | 0.882 | 0.354 | 0.584 | 0.39 | 0.014 | 0.911 | -0.08 | 3.688 | 0.127 | -0.93 | 1.803 | 0.250 | -0.77 | 1.755 | 0.256 | -0.79 |
| | resin available NH4 | 5.041 | 0.041 | 22.681 | 1.99E-06 | 73.93 | 4 | 0.750 | 0.042 | 0.848 | 0.11 | 14.279 | 0.019 | -1.41 | 3.016 | 0.157 | -0.23 | 1.314 | 0.316 | 0.43 | - | - | - |
| | resin available NO3 | 0.022 | 0.885 | 18.468 | 7.64E-06 | 193.43 | 4 | 0.646 | 0.474 | 0.529 | -0.16 | 0.110 | 0.757 | -0.19 | 0.257 | 0.639 | 0.33 | 0.655 | 0.464 | -0.52 | - | - | - |
| | resin available P | 2.313 | 0.151 | 59.156 | 2.20E-09 | 819.12 | 4 | 0.858 | 3.294 | 0.144 | 0.79 | 8.555 | 0.043 | -1.30 | 0.189 | 0.686 | 0.23 | 5.216 | 0.084 | 0.96 | - | - | - |
| | total extractable NH4 | 0.485 | 0.494 | 57.701 | 3.40E-11 | 841.28 | 5 | 0.858 | 13.152 | 0.022 | 1.49 | 0.004 | 0.952 | 0.02 | 1.701 | 0.262 | -0.79 | 0.533 | 0.506 | -0.47 | 5.052 | 0.088 | 1.18 |
| | total extractabel NO3 | 3.286 | 0.086 | 20.726 | 2.74E-07 | 621.46 | 5 | 0.686 | 3.518 | 0.034 | 1.05 | 19.002 | 0.012 | -1.59 | 0.073 | 0.800 | -0.17 | 0.920 | 0.392 | -0.61 | 0.087 | 0.783 | 0.20 |
| | pH | 1.716 | 0.206 | 28.589 | 1.84E-08 | 70354.39 | 5 | 0.747 | 0.281 | 0.624 | -0.35 | 2.761 | 0.172 | 0.85 | 0.057 | 0.823 | -0.16 | 0.655 | 0.464 | -0.52 | 1.933 | 0.237 | -0.84 |
| | %SOM | 5.843 | 0.026 | 180.245 | 6.66E-16 | 3980.05 | 5 | 0.948 | 0.828 | 0.414 | -0.131 | 26.704 | 0.007 | -1.664 | 4.519 | 0.101 | 0.859 | 0.069 | 0.805 | 0.176 | 2.290 | 0.205 | -0.669 |
| Fungal Diversity | Rhizophilic AMF q1 (Shannon exp) | 0.186 | 0.671 | 1.698 | 0.181 | 133.45 | 5 | 0.146 | 0.195 | 0.681 | -0.293 | 4.632 | 0.098 | 0.514 | 0.751 | 0.435 | 0.556 | 0.165 | 0.705 | -0.230 | 0.023 | 0.887 | 0.101 |
| | Edaphophilic AMF q1 (Shannon exp) | 4.611 | 0.045 | 2.313 | 0.082 | 218.51 | 5 | 0.247 | 1.102 | 0.353 | 0.660 | 1.951 | 0.235 | 0.545 | 0.223 | 0.661 | 0.218 | 12.397 | 0.024 | 1.251 | 0.217 | 0.666 | 0.304 |
| | All AMF q1 (Shannon exp) | 5.48 | 0.03 | 1.524 | 0.227 | 474.08 | 5 | 0.211 | 0.146 | 0.722 | 0.254 | 28.222 | 0.006 | 1.540 | 0.281 | 0.624 | 0.350 | 0.402 | 0.560 | 0.415 | 0.161 | 0.708 | 0.267 |
| | All AMF q0 (Richness) | 4.10 | 0.06 | 10.225 | 5.63E-05 | 1226.69 | 5 | 0.530 | 0.259 | 0.637 | -0.323 | 13.997 | 0.020 | 1.514 | 0.751 | 0.435 | 0.556 | 1.616 | 0.273 | 0.778 | 0.157 | 0.712 | 0.192 |
| | Rhizophilic AMF q0 (Richness) | 0.744 | 0.399 | 5.915 | 0.002 | 734.33 | 5 | 0.404 | 0.454 | 0.538 | -0.439 | 2.667 | 0.178 | 0.618 | 0.298 | 0.614 | 0.360 | 0.000 | 1.000 | 0.000 | 0.000 | 1.000 | 0.000 |
| | Edaphophilic AMF q0 (Richness) | 0.212 | 0.650 | 4.092 | 0.010 | 311.45 | 5 | 0.323 | 0.783 | 0.426 | -0.269 | 10.286 | 0.033 | 0.757 | 0.078 | 0.794 | 0.186 | 1.882 | 0.242 | -0.572 | 0.395 | 0.564 | -0.412 |

| Response | AMF | | | | | Roots | | | | | Greenness | | | | |
|---|---|---|---|---|---|---|---|---|---|---|---|---|---|---|---|
| Model statistics | Int | Df | R2 | | | Int | Df | R2 | | | Int | Df | R2 | | |
| | 101.76 | 6 | 0.8 | | | 180.69 | 18 | 0.62 | | | 1883.01 | 10 | 0.82 | | |
| Effect | F | p | *B* | Treatment interaction (F) | Treatment interaction (p) | F | p | *B* | Treatment interaction (F) | Treatment interaction (p) | F | p | *B* | Treatment interaction (F) | Treatment interaction (p) |
| AMF productivity | | | | | | 52.336 | <.0001 | -0.93 | 8.446 | 0.010 | 55.931 | <.0001 | -0.21 | 6.615 | 0.028 |
| Root productivity | 128.804 | <.0001 | -0.24 | 2.281 | 0.182 | | | | | | - | - | - | - | - |
| Greenness (NDVI) | - | - | - | - | - | - | - | - | - | - | | | | | |
| %SOM | 1.307 | 0.283 | -0.33 | - | - | - | - | - | - | - | - | - | - | - | - |
| total extractable NH4 | 1.833 | 0.225 | 0.25 | 1.711 | 0.239 | 0.342 | 0.567 | 0.45 | 10.028 | 0.006 | - | - | - | - | - |
| total extractabel NO3 | 0.697 | 0.436 | -0.31 | 0.569 | 0.479 | - | - | - | - | - | - | - | - | - | - |
| resin available NH4 | 55.541 | <.0001 | 0.95 | 0.009 | 0.929 | - | - | - | - | - | 22.314 | 0.000 | -1.13 | - | - |
| resin available NO3 | - | - | - | - | - | 12.691 | 0.002 | 0.84 | - | - | 69.227 | <.0001 | 0.62 | 0.301 | 0.596 |
| resin available P | 4.397 | 0.081 | 0.4 | 13.872 | 0.005 | - | - | - | - | - | 10.047 | 0.010 | 0.55 | 2.417 | 0.151 |

| | dbRDA Results | | | | | | Post-hoc tests | | | | | | | | | |
|---|---|---|---|---|---|---|---|---|---|---|---|---|---|---|---|---|
| | Treatment | | Time | | | | Under Snow | | Post-Snowmelt | | Late Spring | | Dry Down | | Monsoon | |
| ~ | F | p | F | p | R2 | Df | F | p | F | p | F | p | F | p | F | p |
| All AMF | 1.534 | 0.023 | 2.992 | 0.002 | 0.18 | 40 | 1.040 | 0.479 | 2.563 | 0.032 | 1.020 | 0.494 | 1.071 | 0.479 | 0.7982 | 0.68 |
| Rhizophilic AMF | 2.146 | 0.023 | 2.023 | 0.001 | 0.11 | 38 | 1.265 | 0.326 | 4.424 | 0.018 | 0.809 | 0.606 | 1.986 | 0.131 | 0.763 | 0.526 |
| Edaphophilic AMF | 2.469 | 0.007 | 5.554 | 0.001 | 0.14 | 38 | 2.494 | 0.017 | 4.456 | 0.042 | 0.759 | 0.509 | 1.286 | 0.289 | 0.380 | 0.711 |
| Ancestral AMF | 1.3459 | 0.242 | 0.537 | 0.764 | 0.01 | 32 | 0.510 | 0.713 | 0.809 | 0.484 | 0.820 | 0.594 | 0.726 | 0.746 | 0.335 | 0.778 |
| Unassigned Glomeromycetes | 0.984 | 0.414 | 2.337 | 0.003 | 0.09 | 39 | 1.210 | 0.336 | 2.891 | 0.062 | 0.839 | 0.567 | 0.931 | 0.492 | 1.073 | 0.372 |

| | **dbRDA Results** | | | | | |
|---|---|---|---|---|---|---|
| | All AMF | | Rhizo | | Edapho | |
| | F | p | F | p | F | p |
| AMF productivity | - | - | - | - | - | - |
| Root productivity | 0.815 | 0.746 | 1.829 | 0.018 | - | - |
| Greenness (NDVI) | - | - | - | - | 2.541 | 0.036 |
| Soil Moisture (VWC) | 6.594 | 0.001 | - | - | 1.260 | 0.239 |
| pH | - | - | - | - | - | - |
| resin available NH4 | 2.038 | 0.008 | 3.104 | 0.001 | 1.755 | 0.095 |
| resin available NO3 | - | - | - | - | - | - |
| resin available P | 2.375 | 0.002 | 1.232 | 0.174 | - | - |
| total extractable NH4 | 1.431 | 0.103 | 3.566 | 0.001 | - | - |
| total extractabel NO3 | 1.303 | 0.145 | - | - | 1.826 | 0.024 |
| Residual DF | 29 | | 30 | | 30 | |
| R2 | 0.18 | | 0.12 | | 0.11 | |
